# DNA Replication Asymmetry and Proteostasis Collapse Dynamics

**DOI:** 10.64898/2025.12.07.692845

**Authors:** Regi Avdiaj

## Abstract

Replication fidelity differs across leading and lagging strands and between early- and late-replicating genomic regions, generating asymmetric mutational inputs across proliferative lineages. How such asymmetries influence long-term proteostasis stability has not been quantitatively evaluated. Here, we develop a stochastic generational model linking replication-derived mutation asymmetry to proteotoxic load dynamics under age-dependent declines in clearance. Mutations are drawn from symmetric (H0) or strand- and timing-asymmetric (H1) Poisson processes, converted into proteotoxic load through a binomial misfolding mechanism, and cleared with linearly declining proteostasis efficiency. Proteostasis collapse is defined as a first-passage event when total load exceeds a capacity threshold. Across homeostatic and near-failure regimes, replication asymmetry modestly increases steady-state proteotoxic load (approximately 1– 3%) and leads to modestly earlier collapse. Under clearance-compromised or stress-test conditions, asymmetry substantially amplifies collapse probability and accelerates failure. These results demonstrate that replication-associated asymmetry acts as a weak perturbation when basal load dominates but becomes a significant modifier of collapse risk when proteostasis capacity is reduced. The framework provides a quantitative basis for evaluating how replication dynamics shape long-term proteostasis stability in proliferative tissues.

## Introduction

DNA replication is not a perfectly symmetric process. Leading and lagging strands differ in polymerase usage, fidelity, and mismatch-repair engagement, producing distinct mutational spectra across strands [1, 2, 3]. Replication timing introduces further heterogeneity between early- and late-replicating genomic regions, with late-replicating domains often exhibiting higher mutation rates and distinct mutational signatures [4, 5, 6]. These combined asymmetries influence the distribution and biochemical nature of mutations inherited by daughter cells across proliferative lineages. Although replication-associated mutational asymmetry is well established, its downstream consequences for long-term proteostasis stability remain quantitatively unresolved.

Proteostasis generally declines with age in proliferative tissues due to reduced chaperone availability, diminished proteasomal and autophagic activity, and increased sensitivity to misfolded or aggregation-prone proteins [7, 8, 9, 10]. Even modest increases in proteotoxic burden may, over many divisions, push cells toward thresholds beyond which sustained proteostasis becomes unlikely. Such threshold-like behaviors have been described in multiple proteostasis pathways, including chaperone saturation, aggregate accumulation, and proteasome impairment [11, 12, 13, 14]. These phenomena naturally lend themselves to first-passage modeling frameworks in which collapse occurs when cumulative load exceeds a system-specific capacity limit [15, 16].

Replication-associated mutations can provide a persistent source of misfolding-prone protein variants, particularly when they destabilize native folds or increase aggregation propensity [17, 12, 18]. When combined with basal proteotoxic inputs and age-related declines in clearance capacity, small differences in mutation input may accumulate across many cycles to alter collapse probabilities. Yet the magnitude, direction, and parameter regimes under which replication-derived asymmetry contributes meaningfully to proteostasis failure remain unclear.

Here, we develop a stochastic generational model linking replication asymmetry to proteotoxic load dynamics. Mutations are generated at each division via symmetric (H0) or strand- and timing-asymmetric (H1) Poisson processes, converted into misfolded load through a binomial misfolding mechanism, and cleared with an age-dependent decline in proteostasis efficiency. Collapse is defined as a first-passage event when total load exceeds a capacity threshold. We use this framework to quantify when, and by how much, replication asymmetry influences expected proteotoxic burden, steady-state load, and population-level collapse risk across homeostatic, near-capacity, and clearance-compromised regimes.

## Methods

### Model overview

We developed a stochastic model to quantify how DNA replication–associated mutational asymmetry influences long-term proteotoxic load and collapse risk in proliferative cell populations, consistent with known asymmetries in replication fidelity and mutational spectra [1, 2, 3, 5]. The model consists of: (i) a mutation-generation process per replication, (ii) conversion of mutations into misfolded proteotoxic load plus a basal contribution, and (iii) age-dependent clearance of accumulated load. Collapse is modeled as a first-passage event when total load exceeds a proteostasis capacity threshold *L*_crit_, in line with coarse-grained proteostasis collapse frameworks [15, 16]. Deterministic mean-field analysis is used only to derive equilibrium expectations; all collapse statistics and survival outcomes arise from the full stochastic process.

All load variables *L, B*, and *Y* are dimensionless abstract measures proportional to proteotoxic burden. This reflects the coarse-grained nature of the model, in which chaperone availability, aggregation propensity, and proteasomal or autophagic capacity are collapsed into a single quantitative variable; only relative differences and collapse times are interpreted [17, 12, 10, 14].

### Mutation generation

At each replication cycle *t*, a cell acquires a random number of mutations *M*_*t*_. Under the symmetric hypothesis (H0), mutations follow a Poisson process,

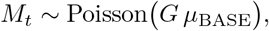

where *G* is an effective genomic size factor and *µ*_BASE_ is a baseline per-unit mutation rate, consistent with treating replication errors as rare, approximately independent events [1, 2].

Under the asymmetric hypothesis (H1), mutations are partitioned into four compartments corresponding to leading/lagging strands and early/late replicating regions, reflecting replication- and timing-dependent variation in mutation rates [4, 5, 6]. For example, the leading–early compartment follows

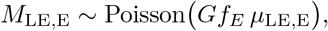

with analogous definitions for

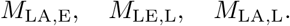

The total number of mutations is

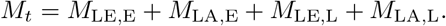

Unless otherwise indicated, we used

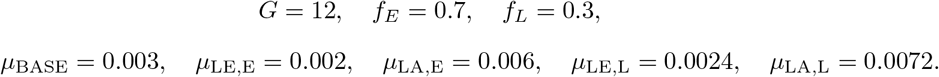

### Misfolding and load accumulation

Each mutation results in production of a protein that misfolds with probability *p*_mis_. The number of misfolded proteins at time *t* is

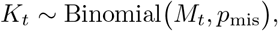

and each misfolded species contributes *Y* units of proteotoxic load. The misfolding-derived increment is therefore

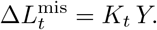

This formulation is consistent with the view that destabilizing mutations and sequence perturbations increase the probability of misfolding and aggregation-prone states, thereby adding to proteotoxic burden [17, 12, 18]. A basal load *B* is added at every cycle to represent all proteotoxic sources not directly linked to replication (e.g. translational errors, oxidative and environmental damage, or constitutive aggregation processes), which are known contributors to age-associated proteostasis stress [14, 18, 10]. The total load increment per cycle is

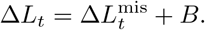

Default values were

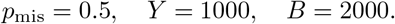

### Proteostasis clearance and age-dependent decline

Total load is updated each cycle via

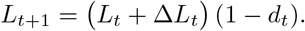

Clearance declines linearly with replicative age:

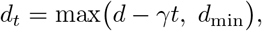

where *d* is the initial clearance fraction, *γ* is the per-division rate of decline, and *d*_min_ prevents clearance from reaching implausibly low values. This captures the experimentally observed decline in chaperone function, proteasomal activity, and autophagic flux with age [ 7, 11, 9, 8, 10].

We used

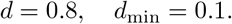

Non-aging simulations used

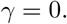

Aging-like simulations used

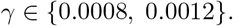

Near-failure regimes (partial collapse by the end of the simulation horizon) used *γ* = 0.0008, whereas stress-test regimes (strongly impaired clearance) used *γ* = 0.0012.

### Mean-field dynamics and equilibrium load

When *γ* = 0, clearance is constant and the expectation of *L*_*t*_ satisfies

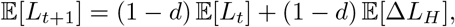

where *H* indicates hypothesis H0 or H1. 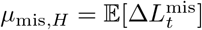. Then

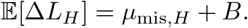

The linear recurrence has the stable fixed point

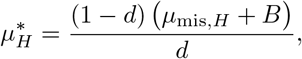

which corresponds to the mean-field steady-state load under constant parameters. Similar linear recurrences and steady-state analyses are used in coarse-grained models of proteostasis capacity and collapse [15, 16].

Deterministic expressions are used only to derive equilibrium expectations; all collapse statistics are obtained from the stochastic simulations.

### Collapse threshold and first-passage criterion

Proteostasis collapse is defined as the first time a lineage satisfies

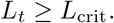

The threshold *L*_crit_ represents a coarse-grained proteostasis capacity limit. This formulation is analogous to threshold-like behaviors described in proteostasis models where overload of chaperone pools, sustained aggregation, or proteasome inhibition produces irreversible functional collapse [11, 13, 16, 12]. Different dynamical regimes were explored by varying *L*_crit_, including (i) near-failure regimes where only a subset of lineages collapse within the simulation horizon and (ii) stress-test regimes where proteostasis is close to immediate failure. Numerical values were chosen to explore dynamical behavior rather than to match empirical datasets.

If a lineage collapsed within the time horizon *T* , the collapse time *T*_coll_ was recorded; otherwise *T*_coll_ = NaN. Survival curves were computed as

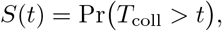

using the empirical fraction of noncollapsed lineages at each time point, as in standard survival and first-passage analyses of proteostasis collapse [16].

### Simulation protocol

Simulations were implemented in Python using NumPy’s random number generator `numpy.random.default_rng`. For each parameter set, multiple independent lineages were simulated in parallel.

Standard parameters included:

- number of lineages: 1000–3000,
- simulation horizon: *T* = 500 for steady-state analyses, and *T* = 2000 for population collapse experiments.

Random seeds were specified to ensure reproducibility. Mutation sanity tests used seeds 42, 111, and 222; population-level experiments used seed 999; and seed robustness analyses repeated the stress-test regime for seeds 0 through 4.

### Validation procedures

#### Mutation sanity checks

We verified that `step mutations` reproduces the expected Poisson means for both H0 and H1. Empirical estimates averaged over 10^5^ draws matched analytic rates, consistent with Poissonian mutation statistics under low per-site error probabilities [1, 2].

#### Single-lineage load dynamics

Load trajectories simulated for *T* = 500 cycles under constant clearance were compared against analytical fixed points 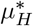. Simulated mean terminal loads matched analytic predictions within a few percent, as expected for linear, weakly fluctuating systems.

#### Basal-load sensitivity

To examine the influence of baseline load *B*, we scaled *B* by factors from 0.25 to 4 and computed percent differences in mean load between H0 and H1. Analytical and stochastic results were consistent: replication asymmetry produces larger relative effects when *B* is small and smaller effect s when *B* dominates, in line with intuition from load–capacity balance models [15, 16].

#### Population collapse near failure

With aging-like clearance decline and a threshold *L*_crit_ adjusted so that only a fraction of lineages collapsed by *T* = 2000, we evaluated collapse fractions, median collapse times, and survival curves. Under these conditions, replication asymmetry modestly advanced collapse.

#### Stress-test regime

Under a lower threshold and more rapid clearance decline, H1 exhibited substantially higher collapse probability and earlier median collapse time than H0, demonstrating amplification of asymmetry effects when proteostasis capacity is severely compromised, consistent with theoretical expectations from models of proteostasis collapse [16].

#### Robustness to random seeds

Stress-test simulations were repeated for seeds 0–4. Collapse fractions and medians were consistent across seeds, indicating that the observed differences between H0 and H1 are robust features of the stochastic dynamics rather than artifacts of a particular random initialization.

### Code availability

All source code, analysis notebooks, and figure-generation scripts are publicly available at: https://github.com/regdough/Replication-Asymmetry-Model

## Results

### Mutation input under symmetric and asymmetric hypotheses

Across 10^5^ independent mutation draws, the symmetric model (H0) produced an average of 𝔼 [*M*_0_] ≈ 0.036, while the asymmetric model (H1) produced 𝔼 [*M*_1_] ≈ 0.101. Thus, H1 generates an approximately 2.8-fold increase in mutation input relative to H0, consistent with the combined effects of strand and timing asymmetry. Both distributions exhibited Poisson-like behavior (variance ≈ mean, maximum observed value = 3).

As a basic validation of the mutation-input module, we compared the per-division mutation counts generated under the symmetric (H0) and asymmetric (H1) hypotheses (Fig. 1). Across 10^5^ simulated divisions, the mean mutation counts were *E*[*M*_0_] ≈ 0.036 for H0 and *E*[*M*_1_] ≈ 0.101 for H1, giving an empirical ratio of ∼ 2.8 that matches the imposed Poisson-rate imbalance. In both cases, the distribution is strongly zero-inflated with a short tail up to three mutations per division, consistent with rare-event Poisson statistics and confirming that the implemented rates are correctly realized in the stochastic simulations.

**Figure 1:**
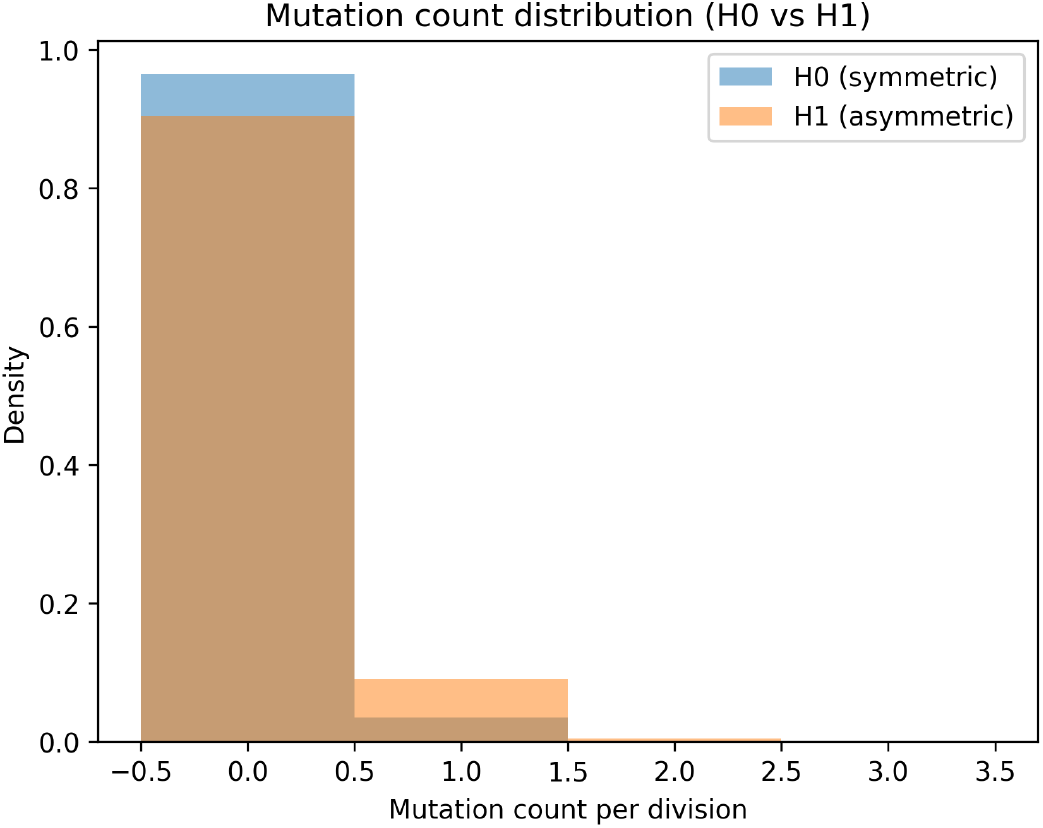
Distribution of mutation counts per division for the symmetric (H0) and asymmetric (H1) models. H1 shows the expected ∼ 2.8-fold increase in mutation input arising from strand and timing biases.

### Single-lineage load dynamics and agreement with mean-field predictions

Under constant clearance (*d* = 0.8, *γ* = 0) and *T* = 500 cycles, simulated terminal loads matched analytical steady-state expectations:

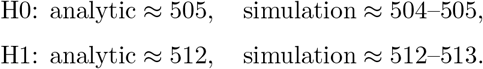

Across 1000–3000 simulated lineages, differences between H1 and H0 remained modest (∼ 1–3%). Stochastic variability around the mean-field limit was small, and no qualitative deviations were observed.

We next examined how asymmetric mutation input alters the final proteotoxic load of single lineages under constant clearance (*d* = 0.8, *γ* = 0). After *T* = 500 divisions, the mean terminal load was 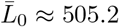 under H0 and 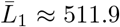 under H1, corresponding to an increase of approximately 1.3–1.6% depending on whether analytic or simulated values are used. The histogram in Fig. 2 shows that both hypotheses produce tightly peaked load distributions with similar variance; H1 is shifted only slightly to the right. This confirms that, under otherwise healthy clearance, replication asymmetry acts as a small perturbation to steady-state proteotoxic load rather than a dominant driver.

**Figure 2:**
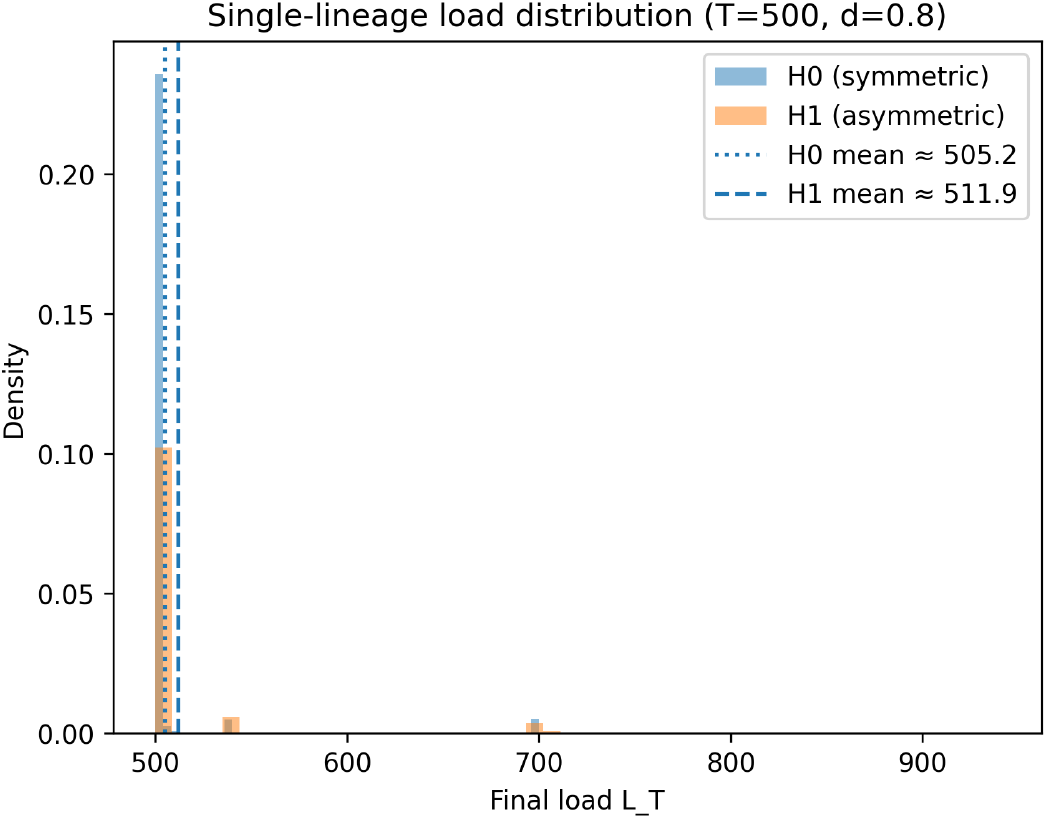
Single-lineage final load distributions after *T* = 500 divisions under constant clearance (*d* = 0.8). H1 exhibits a modestly higher mean load than H0, consistent with the predicted ∼1–3% increase.

### Basal-load sensitivity

Scaling the basal load *B* from 0.25 × to 4× revealed systematic dependence of H1–H0 differences on *B*. When *B* was small, asymmetry contributed 6– 8% relative load increases (analytic and simulated values consistent). As *B* increased, the influence of asymmetry diminished to ∼ 0.4–0.5%.

Representative values were:

| B factor | $\Delta\%$ analytic | $\Delta\%$ simulation |
| --- | --- | --- |
| 0.25 | $\sim 6.3$ | $\sim 8.1$ |
| 0.50 | $\sim 3.2$ | $\sim 4.1$ |
| 1.00 | $\sim 1.6$ | $\sim 2.1$ |
| 2.00 | $\sim 0.8$ | $\sim 1.0$ |
| 4.00 | $\sim 0.4$ | $\sim 0.5$ |

Because *p*_mis_ and *Y* were held fixed, these comparisons isolate the effect of mutation asymmetry itself rather than folding propensity. Changing these parameters would rescale total load but is not expected to alter the qualitative monotonic dependence on *B*.

To quantify how strongly basal proteotoxic input masks the effect of replication asymmetry, we scaled the basal term *B* from 0.25 to 4 times its reference value and computed the percent difference in mean load between H1 and H0.

Analytic predictions and stochastic simulations agreed closely across all *B* values (Fig. 3). For low basal load (*B* factor 0.25), the asymmetric model increased mean load by roughly 6–8%, whereas at the reference value (*B* factor 1.0) the effect was ∼ 1.6–2.1%. At high basal load (*B* factors 2–4), the relative contribution of asymmetry dropped below ∼ 1%. These trends indicate that when background proteotoxic stress is large, replication-derived asymmetry becomes a comparatively weak modifier of total load.

**Figure 3:**
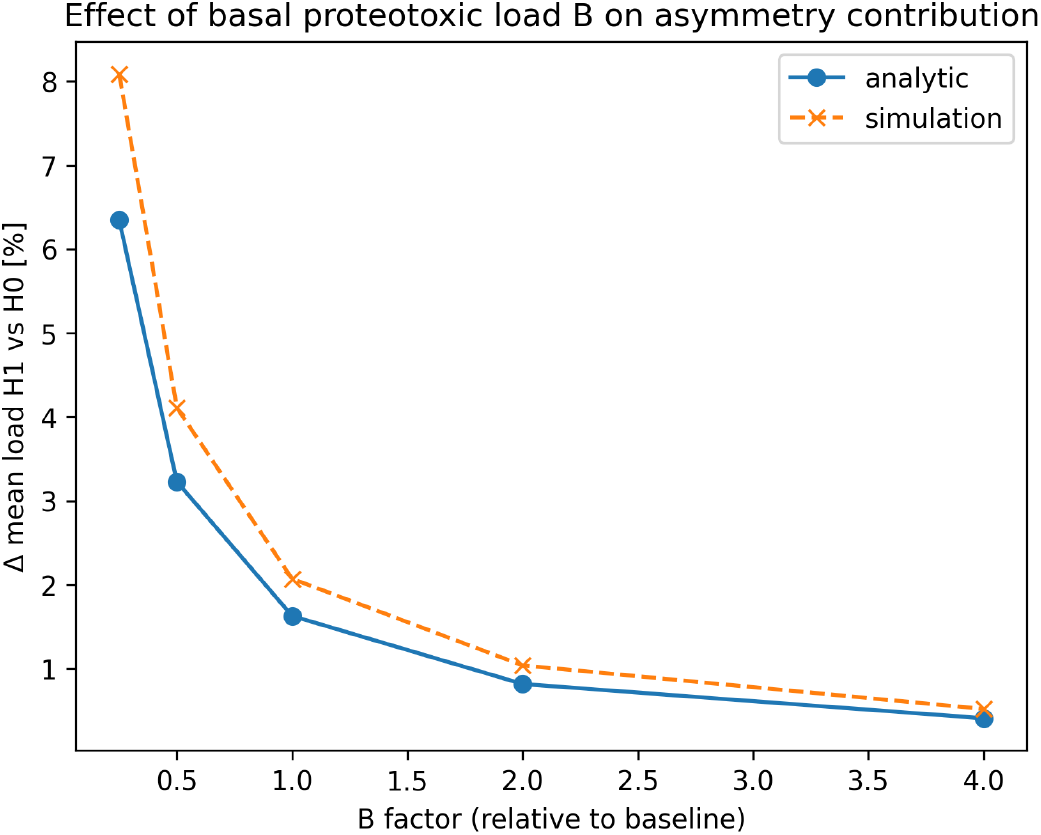
Effect of basal proteotoxic load *B* on the relative contribution of asymmetric mutation input. Both analytic and simulated percent differences decrease as *B* increases, indicating that basal load dominates overall proteotoxic burden in high-*B* regimes.

### Population collapse near proteostasis capacity

To evaluate collapse in marginal regimes, we varied *L*_crit_ across a range of values.

**High thresholds** (≥ 5× 10^4^). No collapses occurred under either hypothesis.

**Moderate threshold** (2 × 10^4^). Differences between H0 and H1 became sub-stantial:

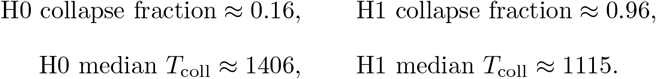

Stochastic spreads in this regime were typically on the order of 100–200 divisions but were smaller than H0–H1 differences.

**Low thresholds** (5 × 10^3^ **and** 1 × 10^4^). Both models collapsed universally, with H1 consistently collapsing 10–30 divisions earlier than H0.

We then probed a near-failure regime by choosing a threshold *L*_crit_ = 2 × 10^4^ such that only a fraction of lineages collapsed by *T* = 2000 divisions. In this setting, the fraction of collapsed lineages by the end of the simulation was low for H0 (∼ 0.16) and high for H1 (∼ 0.96), and median collapse times among collapsed lineages were approximately 1406 divisions for H0 and 1115 divisions for H1. The survival curves in Fig. 4 show that most H0 lineages remained below threshold throughout the time window, whereas H1 lineages exhibited a gradual but pronounced loss of survivors after ∼ 900 divisions. Thus, under moderately stressed proteostasis, replication asymmetry modestly advances collapse and substantially increases the fraction of lineages that fail by a fixed age.

**Figure 4:**
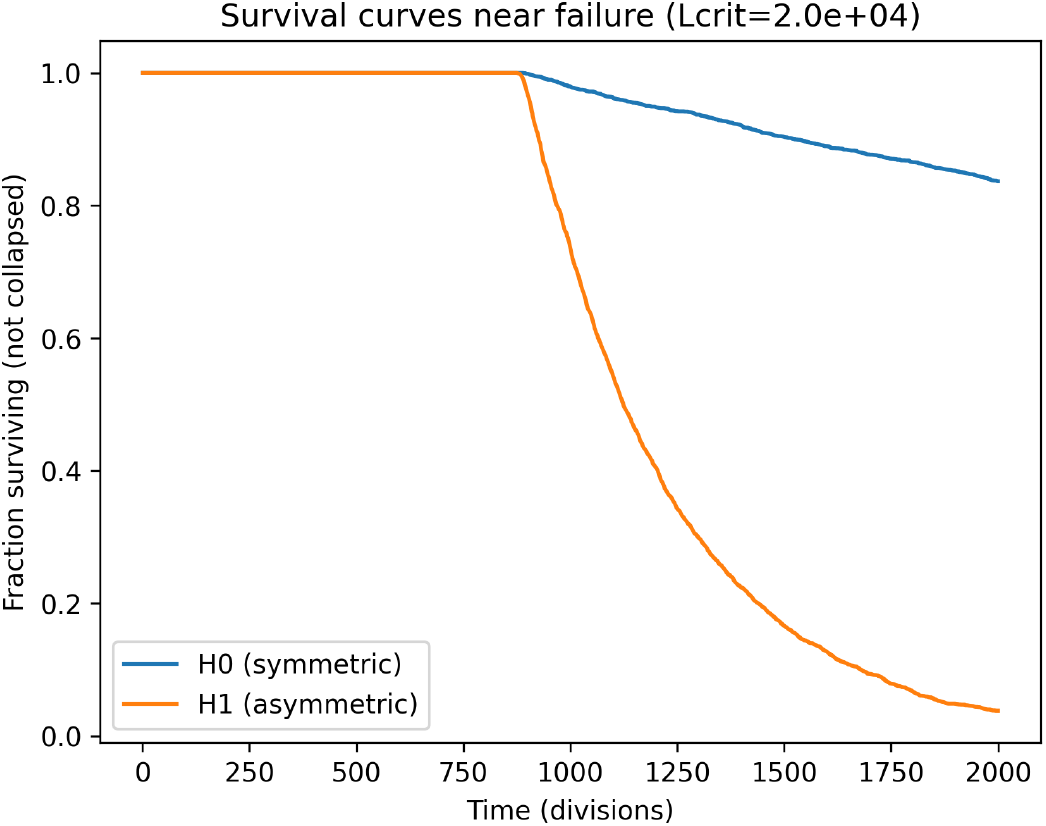
Survival curves for symmetric (H0) and asymmetric (H1) models near the proteostasis capacity threshold (*L*_crit_ = 2 × 10^4^). Asymmetry produces modestly earlier collapse while keeping overall collapse fractions partially over-lapping.

### Stress-test regime: amplified collapse under compromised clearance

Under accelerated clearance decline and reduced proteostasis capacity, asymmetry produced large shifts in collapse behavior:

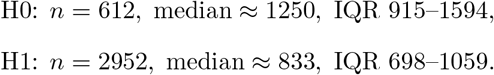

Collapse fractions were:

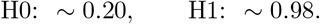

Thus, under severely compromised proteostasis, H1 lineages experienced earlier collapse by approximately 400–500 divisions, and nearly all lineages failed.

Because the asymmetry strength was fixed, these results represent one biologically plausible imbalance. Increasing or decreasing the *µ*-ratios would respectively strengthen or weaken the effect size but would not change the qualitative amplification under stressed clearance.

Finally, we examined a stress-test regime with faster age-dependent decline in clearance (*γ* = 0.0012) at the same threshold *L*_crit_ = 2 × 10^4^. In this regime, collapse was rare for H0 but nearly ubiquitous for H1: the fraction of collapsed lineages by *T* = 2000 was ∼ 0.20 for H0 and ∼ 0.98 for H1, with median collapse times around 1249 and 833 divisions, respectively. The survival curves in Fig. 5 show a slow, shallow decline in H0 survival contrasted with a steep loss of H1 lineages shortly after the onset of clearance impairment. Together, these results demonstrate that replication-derived asymmetry becomes a strong amplifier of collapse risk once proteostasis capacity is substantially compromised.

**Figure 5:**
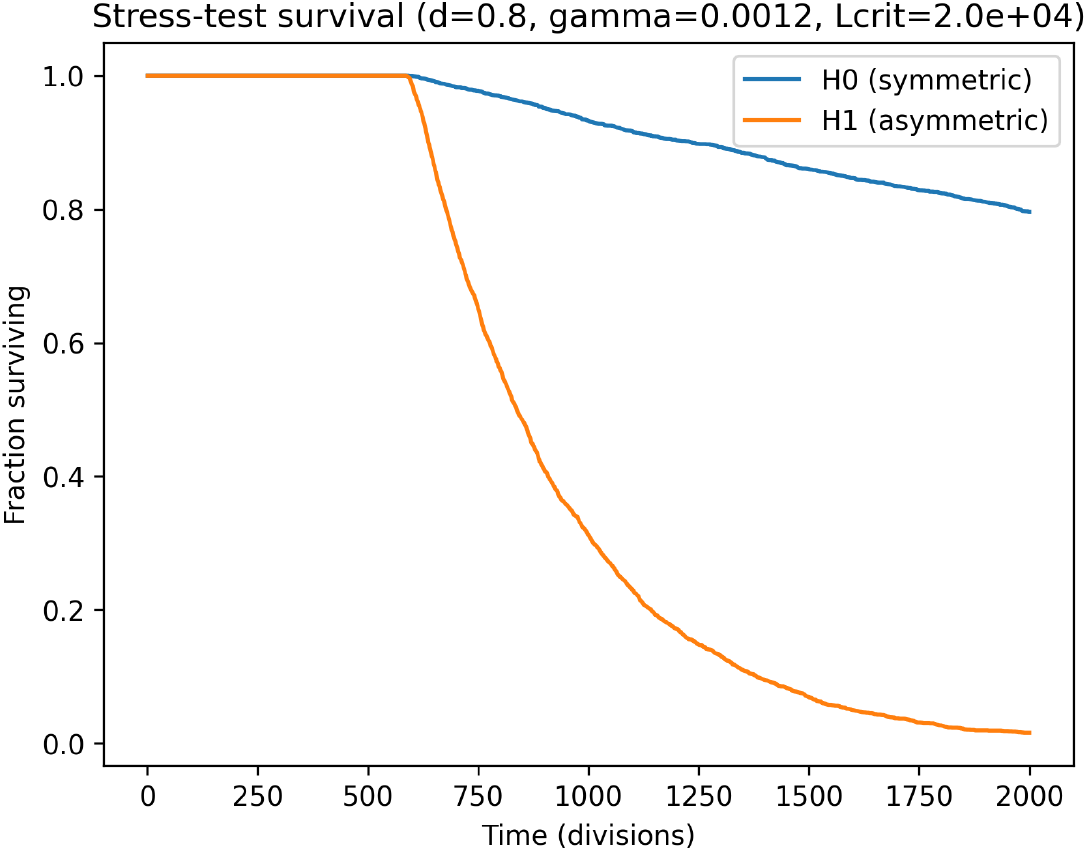
Stress-test survival curves under impaired clearance (*d* = 0.8, *γ* = 0.0012, *L*_crit_ = 2 × 10^4^). Under these conditions, replication-derived asymmetry strongly amplifies collapse probability and accelerates failure.

### Seed robustness

Across seeds 0–4, results were highly consistent:

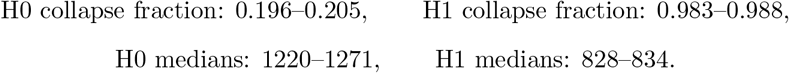

This indicates that the observed differences arise from model structure rather than stochastic initialization.

### Summary of dynamical regimes

Across all simulated regimes:

- Under homeostatic conditions, asymmetry produces small (1–3%) load differences.
- Under near-failure conditions, asymmetry yields moderately earlier collapse and divergent collapse fractions.
- Under severely compromised clearance, asymmetry produces large shifts in collapse timing and nearly complete population failure.

Overall, replication asymmetry remains a weak perturbation when clearance is strong and basal load dominates, but becomes a major modifier of collapse dynamics as proteostasis capacity approaches its limits.

## Discussion

The stochastic framework developed here links replication-derived mutational asymmetry to long-term proteotoxic load dynamics in proliferative lineages. Across a wide parameter range, the model demonstrates that asymmetry produces small but consistent increases in steady-state proteotoxic burden under homeostatic and near-capacity conditions, with typical differences of approximately 1–3%. These modest effects arise because basal load *B* dominates total proteotoxic input, causing mutation-derived contributions to represent only a small fraction of cumulative stress. Under such conditions, even multi-compartment asymmetric mutation inputs contribute only weakly to long-term load.

When proteostasis becomes compromised—either through age-dependent decline in clearance or through a reduced collapse threshold *L*_crit_—the dynamics differ substantially. In these regimes, small differences in mutation-derived input accumulate over many divisions and interact with declining clearance efficiency. As a consequence, collapse probability increases markedly for the asymmetric model, and median collapse times shift earlier by several hundred divisions. These behaviors reflect the threshold-like nature of proteostasis failure: once clearance decreases sufficiently, incremental increases in load can produce disproportionately large effects on survival outcomes.

Sensitivity analyses further clarify the conditions under which replication asymmetry has a measurable impact. Increasing the baseline proteotoxic load *B* proportionally reduces the relative contribution of mutation-derived load, thereby attenuating the difference between H0 and H1. Conversely, decreasing *B* or lowering proteostasis capacity amplifies asymmetry contributions. These monotonic trends, observed analytically and in simulation, suggest that the qualitative conclusions are robust to the choice of basal parameters. Moreover, because the asymmetry strength was fixed to one plausible configuration of strand- and timing-biased mutation rates, the reported effects should be interpreted as one representative case. Increasing or decreasing the *µ*-ratios would scale effect sizes accordingly without altering the qualitative relationships identified here. Importantly, the numeric values used here are not intended to predict collapse ages for specific tissues; they serve as illustrative parameters to explore how asymmetry interacts with declining proteostasis capacity at the level of dynamical regimes.

Stochastic variability also plays an important role. Even when differences in expected load are small, first-passage events are sensitive to fluctuations, especially under declining clearance. Interquartile ranges for collapse times remained broad across both near-failure and stress-test regimes, and seed-robustness analyses showed consistent collapse fractions and medians across replicate simulations. These observations confirm that the separation between H0 and H1 is not an artifact of specific random seeds, but instead emerges from the structure of the generational dynamics.

Several simplifications are inherent to the model. Proteotoxic load is expressed as a single abstract quantity, subsuming diverse molecular processes such as chaperone binding, aggregation, and degradation. Mutation effects are treated uniformly through a fixed misfolding probability, and the clearance decline is represented by a linear function of replicative age. These abstractions enable tractable analysis but do not capture the full biochemical complexity of proteostasis networks. Nonetheless, the model provides a quantitative baseline for evaluating how replication-derived asymmetry interacts with declining clearance to influence collapse risk.

Overall, the results indicate that replication-associated asymmetry acts as a weak perturbation under strong homeostatic control but becomes a meaningful modifier of collapse probability when clearance efficiency deteriorates. This distinction highlights the importance of proteostasis capacity in shaping the long-term consequences of mutation-input asymmetries. The framework establishes a foundation for future theoretical work on how genomic replication dynamics influence cellular aging trajectories and the emergence of failure thresholds in proliferative tissues.

## Conclusion

This work develops a stochastic generational framework to examine how replication-derived mutational asymmetry may influence proteotoxic load dynamics under age-dependent declines in clearance. Across homeostatic and near-capacity regimes, the model generates small but consistent differences in steady-state load between symmetric and asymmetric mutation inputs, reflecting the predominance of basal proteotoxic sources over mutation-derived contributions. Under reduced proteostasis capacity—via declining clearance or lower collapse thresholds—these modest differences can accumulate across many divisions and lead to earlier and more frequent collapse events. These qualitative behaviors are robust across variation in basal load, threshold placement, and random initialization, but should be interpreted as theoretical regime analyses rather than predictions for specific biological systems.

The framework suggests that replication asymmetry functions as a weak perturbation when clearance is efficient but can act as a meaningful modifier of collapse probability under compromised proteostasis conditions. Given its simplified representation of mutation effects and proteostasis pathways, the model serves as a baseline for theory-driven exploration of how mutation-input asymmetries interact with declining capacity. Future extensions may incorporate heterogeneous mutation impacts, nonlinear clearance dynamics, or system-specific asymmetry parameters to broaden the space of theoretical regimes examined.

